# Bipartite structural evaluation: Extended network generation model and corrected randomization techniques

**DOI:** 10.1101/2021.10.21.465267

**Authors:** Harry de los Ríos, María J. Palazzi, Aniello Lampo, Albert Solé-Ribalta, Javier Borge-Holthoefer

## Abstract

Understanding the structural organization of bipartite networks is essential for analyzing complex systems across diverse domains, including ecological communities. Such networks often display distinct architectural patterns—particularly nestedness and modularity—that influence system stability, diversity, and dynamics. Despite extensive research, accurately replicating these structures remains challenging, partly due to limitations in existing null models and synthetic network generation methods. Traditionally, nested and modular network ensembles are derived either from *ad hoc* constructions or large sets of random networks, with structural patterns evaluated *a posteriori*. In this article, we present a comprehensive framework that integrates advanced null models with a synthetic network generator to rigorously assess bipartite network architectures. We evaluate the statistical significance of nestedness, modularity, and in-block nestedness scores using five null models across approximately 25,000 synthetic bipartite networks. Our analysis identifies systematic biases in certain models and introduces a Corrected Probabilistic model to address these issues. By combining the analytical simplicity of our network generator with a robust null model, our framework enhances the analysis of bipartite networks, offering valuable tools for ecology and beyond.

## I. INTRODUCTION

Over twenty years ago, research in community ecology was deeply challenged by the evidence that biota interactions, rather than being randomly assembled, manifest clear architectural patterns. In 2003, the pioneer paper of Bascompte *et al*. [1] showed that a significant amount of mutualistic plant-animal networks are organised according to well-defined nested arrangements. Along the same line, Olesen *et al*. [2] unveiled the prevalence of modular configuration among pollination communities with a high (*>* 150) number of species. The observation of these patterns in real ecological networks fuelled a long series of works aimed at establishing how the organisation of species interactions affects the resulting community dynamics: nestedness was recognised to be linked to the promotion of diversity [3], the maximisation of abundances [4] and feasibility [5–7]. On the other hand, modularity was associated to the improvement of local asymptotic stability [8, 9]. These results led to a great consensus about the crucial role of structural arrangements in the behaviour of natural systems [10–14]. Arising from these findings, many methodological advances and tools emerged, to measure and identify those interaction patterns in an automatic and efficient way [15–21].

One would think that these theoretical and methodological achievements rest on well-developed tools to assess their statistical validity. This would imply both the capability to generate large ensembles of networks with prescribed structural features, and also solid procedures to randomize networks to disentangle genuine emergent properties from mere by-products. However, many studies have so far addressed these problems with *ad hoc* strategies: the former, by testing the presence of a given pattern over networks with a random organization [6, 7, 10, 13], or by controlling the structure in a contrived way [9, 12]. The latter, by shuffling network edges with methods that may render biased results.

Another pressing issue relates to the structural arrangements under consideration. In addition to nestedness and modularity, there are other patterns worth being considered, *e*.*g*., in-block nestedness, which describes networks made up from weakly interlinked blocks with internal nested organization [22–24]. This pattern was hypothesized by Lewinsohn in 2006 [22], and it has received much attention in the last few years [23, 25] due to its predicted emergence from micro-[26, 27] and macroscopic [28] mechanisms, and also because of its identification in real mutualistic communities [29]. If the production of synthetic ensembles of structured bipartite networks was feebly addressed for nested or modular structures, it is close to inexistent for hybrid in-block nested arrangements (but see [30]).

In this work, we fill these methodological gaps by, first, introducing BUNGen (Bipartite and Unipartite Network Generator), an integrated synthetic benchmark that generates unior bipartite networks, with varying levels of prescribed planted structures with few and simple parameters. The model we present is a generalization of the model proposed in [24] so as to construct networks with a specified number of blocks (with equal or different sizes) and connectance, inducing different structural signatures. The model is defined by an exact mathematical recipe, remindful in some aspects to that of Beckett and Williams [31], although the latter does not include any functionality to generate modules nor hybrid structures. In developing an analytically-grounded generative model, our contribution aligns with current trends in the structural analysis of networks, which involve utilising Stochastic Block Models and Bayesian inference [32]. Their aim is to decompose networks into blocks of nodes with shared network characteristics, and to determine independent edge probabilities based solely on group membership. In general, these tools require models be defined in a probabilistic framework to compute the likelihood of each model given the observed data –and this is what our model delivers.

Then, on top of BUNGen, we analyze the statistical significance –in terms of *z*-scores– of three rich sets of bipartite matrices with nested, modular and in-block nested configuration. We consider four different null models from the literature, and detect the presence of ambiguities in assessing the various network descriptors, corresponding to over- (or under-) estimations of the given arrangements. To overcome the observed inconsistencies, we introduce an alternative approach, the Corrected Probabilistic model, that helps to reduce the observed under- and overestimation, and highlights the need for the development of new frameworks that take into account those biases. These contributions are wrapped into a single compact framework, consisting of software repositories that enable the generation of the synthetic networks with the specified planted structures, the calculation of nestedness under different formulations, optimization algorithms for the maximization of modularity and in-block nestedness, and the construction of the reviewed null models.

The remainder of the article is organised as follows: Section II introduces the formulation of our synthetic network generation model. Section III starts with a brief recapitulation of some representative randomization schemes for bipartite networks, highlighting their pros and cons. Then, we introduce our reshuffling strategy, which retains some desirable features while diminishing the disadvantages of those reference models. Finally, Section V discusses the main open challenges for further developments.

## II. SYNTHETIC BENCHMARK

In this section we present the analytical foundations of the generative model aimed to cover the variety of structures introduced so far, namely, to build networks with planted nested, modular, and in-block nested arrangements, as well as completely random ones. As a reminder, a description of nestedness, modularity, and in-block nestedness is available in the Supplemental Material (Section S1). For convenience, the model is available as an open-source package. The formulation below corresponds to the case of bipartite networks (*e*.*g*., *N* animal and *M* plant species), and the reduction to unipartite ones is achieved by simply considering *N* = *M*.

The generation of an *N* × *M* adjacency matrix with a given pattern depends first on the number of blocks *B* ≥ 1 with row and column numbers provided respectively by {*r*_*α*_} and {*c*_*α*_} (*α* = 1, …, *B*). Note that 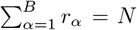 and 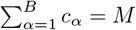.

The intra-block structure is obtained by first generating a perfect nested configuration, whose shape depends on the parameter *ξ* as

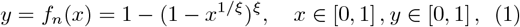

inspired by the unit ball equation. In this equation, *ξ* governs the shape of the separatrix –ranging from convex (0 *< ξ <* 1), to diagonal (*ξ* = 1), to concave (*ξ >* 1), see top row Figure 1.

**FIG. 1.**
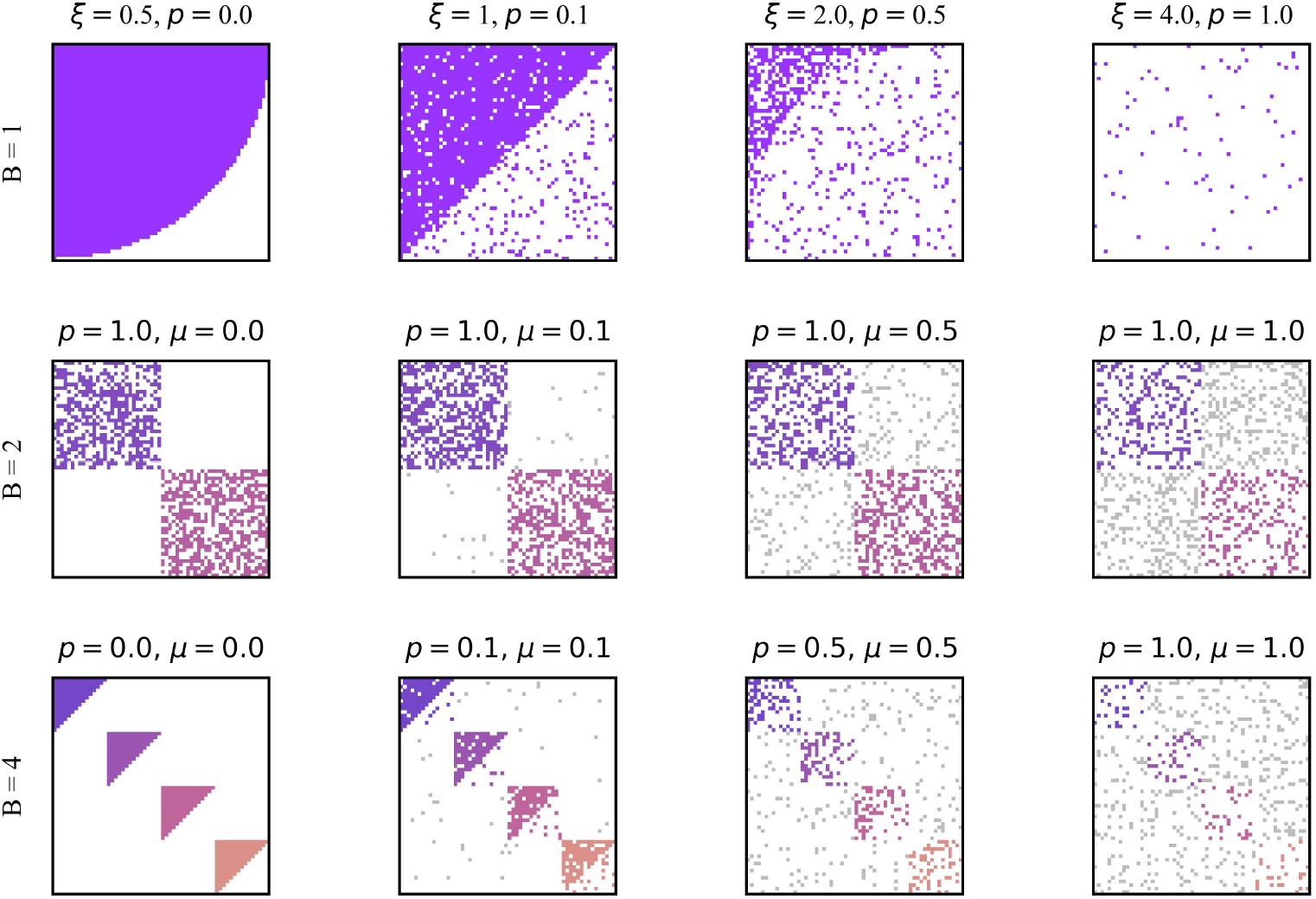
Examples of synthetic network generation with the model introduced here. The top row shows the results for the generation of nested networks (*B* = 1) and the effects of varying the intra-block noise *p* parameter. Middle row exhibits results for two-module networks with varying levels of *p* and the inter-block parameter *µ*. Finally, the bottom row corresponds to in-block nested networks, and again provides some examples of the effect of the noise parameters *p* and *µ*.

Specifically, the *α*^*th*^ block presenting *r*_*α*_ rows and *c*_*α*_ columns is constructed by tessellating the unit ball domain in Eq. 1 into *r*_*α*_ × *c*_*α*_ cells, and adding a link into each entry whose centre lies above the curve. Note that the parameter *ξ*, defining the slimness of the block-nested profile, fixes the number of links *E* and so uniquely determines the density of the network (or connectance, in the network ecology jargon) *d* = *E/*(*N* × *M*).

As mentioned, perfectly nested blocks are rarely found in real systems. Thus, we introduce the concept of noise in the model, relying on a dual-step procedure: a fraction *p* of links lying above the curve in Eq. 1 are randomly selected and then distributed across the empty entries of the block. The top row of Figure 1 illustrates the effect of increasing *p* levels (from left to right), towards a completely random setting (*p* = 1). Note that nested matrices are obtained by setting *B* = 1, *i*.*e*., considering only one block, and are fully characterised by the choice of *ξ* and *p*.

For compartmentalised networks, we introduce inter-block perturbations with an additional noise procedure ruled by the parameter *µ*: for each block, each link is removed with probability

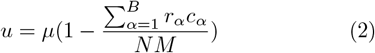

and then reassigned uniformly at random to connect a node of the original block to another belonging to a different one. Note that the term in parenthesis simply reflects the probability of choosing an edge outside the block-diagonal structure. The effects of *µ* are illustrated in the second and third rows of Figure 1, with increasing levels of inter-block noise from left to right. Note that purely modular networks (second row) are obtained setting *p* = 1, *i*.*e*., the internal structure of blocks is assumed to be random.

The generative process outlined above can be analytically described in terms of edge probabilities, simplifying the process of algorithmic design. For the sake of clarity, we can split the probability of having a link between nodes *i* and *j* for the cases in which *i* and *j* belong to the same (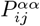) or different (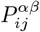) blocks:

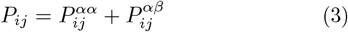

The probability of placing an intra-block link is defined as:

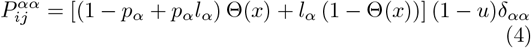

where *p*_*α*_ is the block *α*-specific “intra” noise, and the term *l*_*α*_ corresponds to

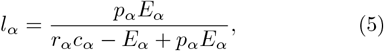

giving the probability of selecting links among the removed ones, with *E*_*α*_ being the number of links in block *α*.

With this in mind, the interpretation of Equation 4 is clear: the term (1 − *p*_*α*_) embodies the probability of not altering a link in the block *α*; the accompanying *p*_*α*_*l*_*α*_ expresses the probability of recovering a link after removal in the random dispersion of removed links in the block *α*; and the Heaviside function Θ(*x*), with *x* = *j/r*_*α*_ − *f*_*n*_(*i/c*_*α*_), restricts the previous processes to the noiseless block portion. On the other hand, *l*_*α*_ (1− Θ(*x*)) describes the alternative probability of a link being dispersed outside the noiseless block portion. Finally, (1 − *u*) ensures that affected links are not those to be spread across blocks (see Eq.2), just as the Kronecker delta *δ*_*αα*_ guarantees that nodes *i* and *j* belong to the same block *α*.

On the other hand, the probability of an inter-block link (involving nodes *i* ∈ *α* and *j* ∈ *β*) is

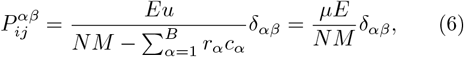

where the numerator takes into account the amount of removed links, while the denominator embodies the possible matrix entries where those links can be placed, *i*.*e*., anywhere outside the defined blocks. The Kronecker delta *δ*_*αβ*_ guarantees that nodes *i* and *j* belong to different blocks.

## III. NULL MODELS

The appropriate selection of a null model has been a center of debate in network ecology for decades [33, 34]. In general, the approaches for the generation of random networks can be classified under three large families, which we summarize below. Following the nomenclature employed in [35], we will refer to them as EE, FF, and PP models.

The Equiprobable-Column or Equiprobable-Row (EE) model represents the least constrained scheme. EE is a probabilistic binary null model that assumes that all the interactions between pairs of nodes –row-to-column links– are equally probable, and that such connection is given with probability 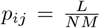, where *L* is the total number of connections in the original matrix. This model generates Erdös-Rényi networks with nodes exhibiting a varying number of connections around a well-defined average degree ⟨*k*⟩, preserving *L* on average. The model tends to falsely detect significant nested patterns, *i*.*e*., it is prone to Type I statistical errors [31, 36, 37].

Fixed Column/Fixed Row (FF) models lie in the other extreme of the spectrum [17], enforcing row-nodes’ and column-nodes’ degrees to be strictly preserved. It is thus the most restrictive null model. The usage of this type of model to assess the probability that an observed pattern could be explained by chance alone has been applied since the 1970s [34]. A typical way for generating random networks under this constraint is through the application of some form of swap algorithm, performing exchanges between pairs of interacting nodes without altering the network’s degree sequence [38].

Noteworthy, generating random matrices by exactly preserving row and column degrees is far from being a trivial task, and so implementations of FF models are quite sophisticated. Swap methods tend to generate repeated matrices that closely resemble the original network. Thus, in order to produce a truly representative set of matrices within the universe of possible matrix configurations, a very large number of matrices need be generated [39], which might not be possible in some scenarios. As a consequence, this model is susceptible to Type II errors, particularly, when it comes to the assessment of nested patterns [35, 37]. On the other hand, it also tends to generate matrices with more checkerboard units, *i*.*e*., 2×2 submatrices containing 1’s in the main diagonal and 0’s outside, or vice versa, than expected by chance [40], producing a biased sample. To solve the latter downside, scholars have introduced variations of the swap methods, and alternative methods that are able to generate a uniform sample of null matrices with fixed row and column degrees [40–44].

In this work, we employ the Curveball algorithm as a reference for FF models. It was introduced by Strona and co-authors [41] as a fast and efficient alternative to generate random matrices with fixed nodes’ degrees, guaranteeing a uniform sample [45].

Row-Proportional/Column-Proportional (PP) models stand somewhere between the extremes –and limitations– described above [17]. They represent an intermediate alternative to generate networks that preserve node degrees on average, thus mitigating the bias towards Type II (FF) or Type I (EE) errors. In these models, interactions between node pairs are defined independently for each individual link on the basis of a Bernoulli trial. From this general principle, there exist different flavors along the loose-strict continuum. We introduce here two variants which, like Curveball, will be used for the sake of comparison against our proposal.

Bascompte’s probabilistic model [1] is perhaps the most popular for the assessment of statistically significant structural patterns (particularly nested ones). In this model, the probability of interaction of a pair of nodes is proportional to the nodes’ degrees of the original network as:

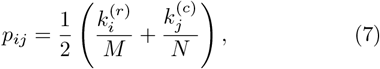

where 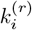 is the degree of the *i*−th row, 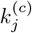 is the degree of the *j* − th column, and *N* and *M* are the number of row and column nodes, respectively.

On the other hand, we consider maximum-entropy exponential random graphs, which were introduced in the last decade [46]. This randomizing framework is based on the maximization of the entropy of a null ensemble, leading to the Exponential Random Graph model [47]. This framework facilitates the analytic computation of statistics over an ensemble of networks with a given set of fixed properties (connectance, average degree of vertices, clustering coefficient, etc.). Under this formulation, the expected values and standard deviation of the network properties are calculated by finding a probability distribution over an ensemble of networks which, on average, preserve the network’s degree sequence. This approach can be applied to both uni- and bipartite networks, and has been recently exploited to assess the significance of nestedness structural arrangements in economic [48], and ecological [49] networks.

Finally, we introduce a new scheme that pivots on Bascompte’s approach. To do so, we revisit Equation 7. This expression can be interpreted as two consecutive sampling processes: one in which we sample the links of each row homogeneously at random, retaining the link only half of the times (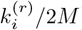), and a second sampling process in which we do the equivalent procedure per columns (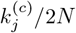). Understanding the process this way, we can reveal two aspects that affect the model performance in maintaining, in average, row and column degree of the original network.

The first aspect regards the oversampling of links per row (and column), and is a consequence of maintaining the two random sampling processes independent. Consider we first distribute links along the rows. Then, when distributing per columns, there are already some links present which need to be considered when sampling the degree of the columns, or else this dual step process will generate more links than expected. We can quantify this excess of degree by computing, on the basis of Eq. 7, the expected degree of a row in the ensemble:

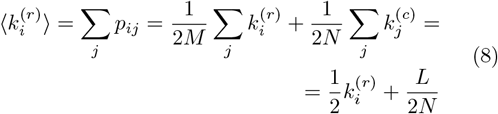

where 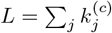 is the number of links of the network. As we can see, the error basically depends on the ratio between *L* and the number of row nodes, which is in essence the average degree. Considering the magnitude of this error and the heavy tail distribution of nodes’ degrees usually observed in ecological networks, a significant overestimation on low degree nodes is expected.

The second aspect that needs to be addressed regards the chance to sampling twice the same link, which will decrease the overall expected connectance of the network and consequently the expected degree of some nodes. The probability of sampling twice a link is equivalent to the probability the adjacency matrix takes weight 2 during the dual step process, *a*_*ij*_ = 2, which can be obtained jointly considering the sampling of the same link by rows, 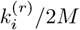, and by columns, 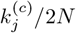:

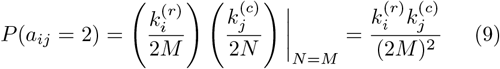

Note that we usually consider a presence/absence adjacency obtained by thresholding the values of the adjacency matrix if a weight is larger than 1. This threshold is implicit in Eq. 7. From Equation 9 we can see that the larger the degree of nodes, the higher the chance of sampling twice the edge that joins them. Then, the larger the under-sampling of its degree on the ensemble. This is particularly problematic in larger degree nodes which turn out to be the most important nodes of the network.

Our Corrected Probabilistic model, inspired by the original model by Bascompte, introduces a variation that alleviates the degree deviations in Eq. 8 and 9. As in the original formulation, the interaction between pairs of species will be estimated considering the nodes’ degrees of the observed interaction matrix. Although we eventually provide analytical expressions for the probability of having an interaction between species, we consider important to describe the process behind these expressions. The Corrected Probabilistic Model is based on two steps: in the first one, we distribute homogeneously at random the interactions of each row among the existing columns of the adjacency matrix; then, in the second step, we proceed to do the same process for each column. However, in the second step, since we already have links placed by the first one, we need to prevent re-sampling links already established, so as to not exceed the expected degree of columns. This process is illustrated in Fig. 2. Probabilistically, the chance of placing a link in between species *i* and *j* in the first step is equivalent to the original model: 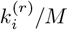. Then, we have to formalize that a link can only be placed if it has not been placed on the first step, that is 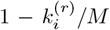 to compensate the column degree induced by the sampling of rows (first step). This induced degree can be easily calculated as 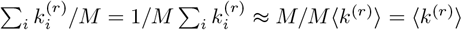, where ⟨*k*^(*r*)^⟩ is the average degree of rows. Thus, the induced degree needs to be discounted when distributing links in the second step, this sampling becomes: 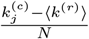. See that, the aforementioned process will exactly maintain in average the degree of rows and will, approximately, maintain the degree of columns. To compensate the induced error between rows an columns, we can assume that half of the times this dual process starts by rows, and the other half by columns. Considering all of the above, we can now obtain the corrected probability of interaction between species *i* and *j* as follows:

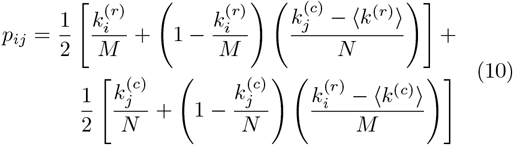

**FIG. 2.**
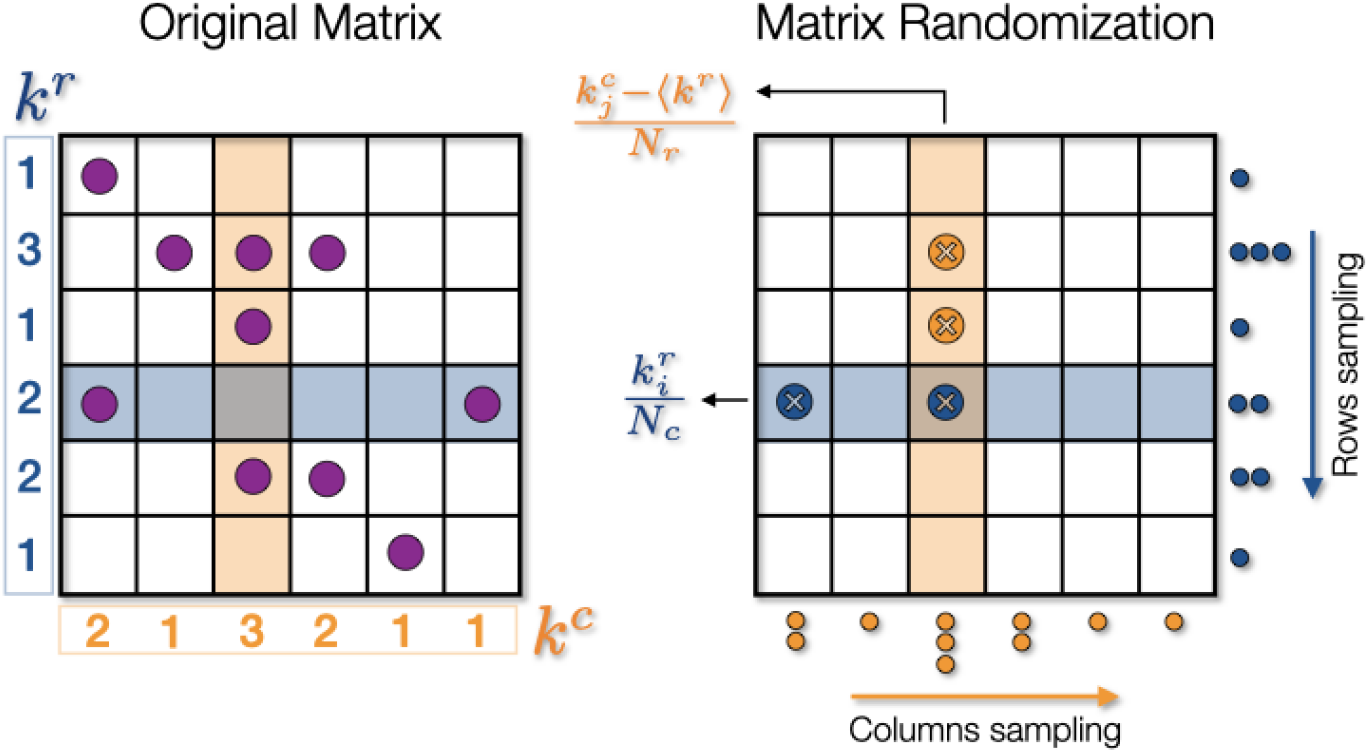
Illustration of the Corrected Probabilistic model. In the left we have the original matrix, while on the right the result of the null model randomisation is shown. In the model, the presence of a link is ruled by Equation 10, which extends Bascompte’s model by taking into account the fact that row and column sampling processes are not independent. Particularly, the figure shows a situation where the row sampling introduces a link (in the fourth row) which affects the third column degree. Accordingly, the probability to have links in a given column is corrected by subtracting a quantity embodying the probability that a fraction of these links already exists because of the row sampling.

The accuracy of the three PP models in nested matrices is provided in Figure 3. Taking a synthetic network with planted nestedness and low noise level (left), top panels show, from left to right, the pairwise interaction probabilities *p*_*ij*_ that result from the maximum entropy, Bascompte’s Probabilistic, and Corrected Probabilistic models. The bottom panels confront expected (actual) degree against the obtained degree after sampling the nested probability matrices (with one standard deviation represented as error bars). Notably, the max-entropy model recovers exactly (with minor fluctuations) the degree sequence of the reference network. Also, it is apparent that, as predicted, lower degree nodes suffer from a significant overestimation in the Probabilistic model, while the opposite happens for the largest degrees. The Corrected Probabilistic model largely solves the inaccuracies of the original model. Still, the main problem of the proposed extension relates to nodes with low degree. This comes from the fact that, although we can compensate the induced degree of the first step into the second, the model is not able to remove links if the induced degree already exceeds the expected degree of columns. Noteworthy, EE and FF models are not included in this figure –their scatter plots would render a trivial picture. The same figures for other network arrangements are available in Fig. S2 of the Supplemental Material. In them, the only evident degree fluctuations stem from the Probabilistic model.

**FIG. 3.**
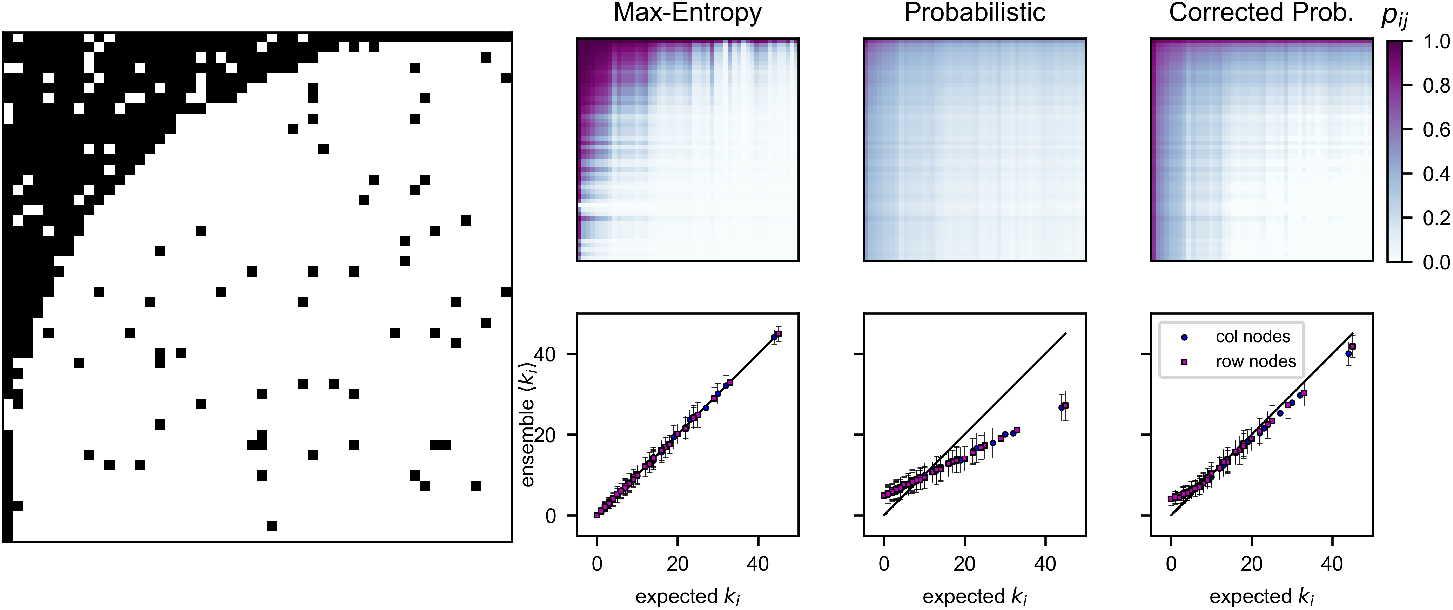
Comparison between a synthetic network (left) with planted nested structure, and their corresponding matrices of interaction probabilities *p*_*ij*_ (top right row) produced by the three PP models: maximum-entropy, Probabilistic and Corrected Probabilistic, respectively, and the scatter plots showing the fit between the obtained average degree sequences with each model and the real degrees of the matrix (bottom right row). Error bars in the scatter plots indicate one standard deviation above and below the average. The same comparisons between the three models for matrices with planted modular and in-block nested structures are available in Fig. S2 of the Supplemental Material.

## IV. RESULTS

In the following, we present the results concerning the test of the five null model classes against the three architectural patterns of interest. For a systematic exploration, Sections IV A and IV B exploit the synthetic generative model introduced in Sec. II. With it, the construction of nested matrices is performed by keeping fixed the number of blocks (*B* = 1) and varying values of *ξ* and *p*, as in the top row of Fig. 1. For the set of purely modular networks, we fix *ξ* = 1.5 and *p* = 1, and smoothly modify *B* and *µ* to cover a wide range of modularity values, *e*.*g*., second row of Fig. 1. Finally, the generation of in-block nested networks is carried out setting *B* = 4, *ξ* = 1.5 and varying levels of *p* and *µ, e*.*g*., third row of Fig. 1. At the end of the process, we generated around 9000 networks for each of the three patterns of interest, *i*.*e*., ~ 2.5 × 10^4^ networks in total. For each generated network, we created 100 random surrogates considering each null model. Among these networks, the number of rows and columns were varied within the range of 30 to 60. Finally, in Section IV C, we perform the statistical significance test on a large collection of empirical networks.

The presence of the arrangements of interest for all these networks –synthetic and empirical– is quantified using specific measures. The level of nestedness is assessed with 𝒩, a NODF-like measure, as defined in [24]. Modularity *Q* was optimized using the extremal optimization algorithm [50]. In the case of in-block nestedness, we employed the optimization method defined in [24].

### A. Resemblance of original *vs*. randomized networks

To understand which kind of networks each null model generates, we start by analyzing to what extent the original network resembles those of the corresponding null model ensembles. Fig. 3 already provided some hints, evaluating the accuracy at maintaining, on average, the degree of the original matrix in PP models.

For a wider look at the outcomes of the null models, we now study the similarity between the random matrices and the original synthetic ones over the range of parameters of the synthetic network generator model introduced in Sec. II. To quantify such resemblance, we employ the Jaccard distance *J*_*d*_, which accounts for the level of dissimilarity between two binary arrays –the original and its random counterparts, in this case. It should be noted that, in the case of binary arrays, *J*_*d*_ = 0 indicates that the matrices are identical.

Top row in Figure 4 shows the average Jaccard distance ⟨*J*_*d*_⟩ over the nested *ξ* − *p* parameter space for each null model scheme. Similar results for the modular and in-block nested matrices are shown in Fig. S3 of the Supplemental Material. As expected, the results for the EE and the probabilistic models show high values of the ⟨*J*_*d*_⟩ over the entire *ξ* − *p* parameter space, confirming that the randomized matrices generated by these models are substantially different from the original one. Turning to the FF (represented by the Curveball algorithm) and Maxentropy models, we observe a wide region of the *ξ* − *p* parameter space in which ⟨*J*_*d*_⟩ drops to intermediate and low values –specially for networks with small noise *p*, regardless of the slimness of the nested structure dictated by *ξ*. These results for FF and Max-entropy confirm that, at least for nested networks, the resulting null matrices are very similar to the original one, since the models only allow for a very little deviation on the degree sequences –none in the FF model. Hence, these models might easily fall into Type II errors. The Corrected Probabilistic model, instead, provides a good trade-off, offering a better fit of the degree sequences, while generating null matrices less similar to the original one, as evinced by the overall intermediate values of the ⟨*J*_*d*_⟩ over the nested *ξ* − *p* parameter space.

**FIG. 4.**
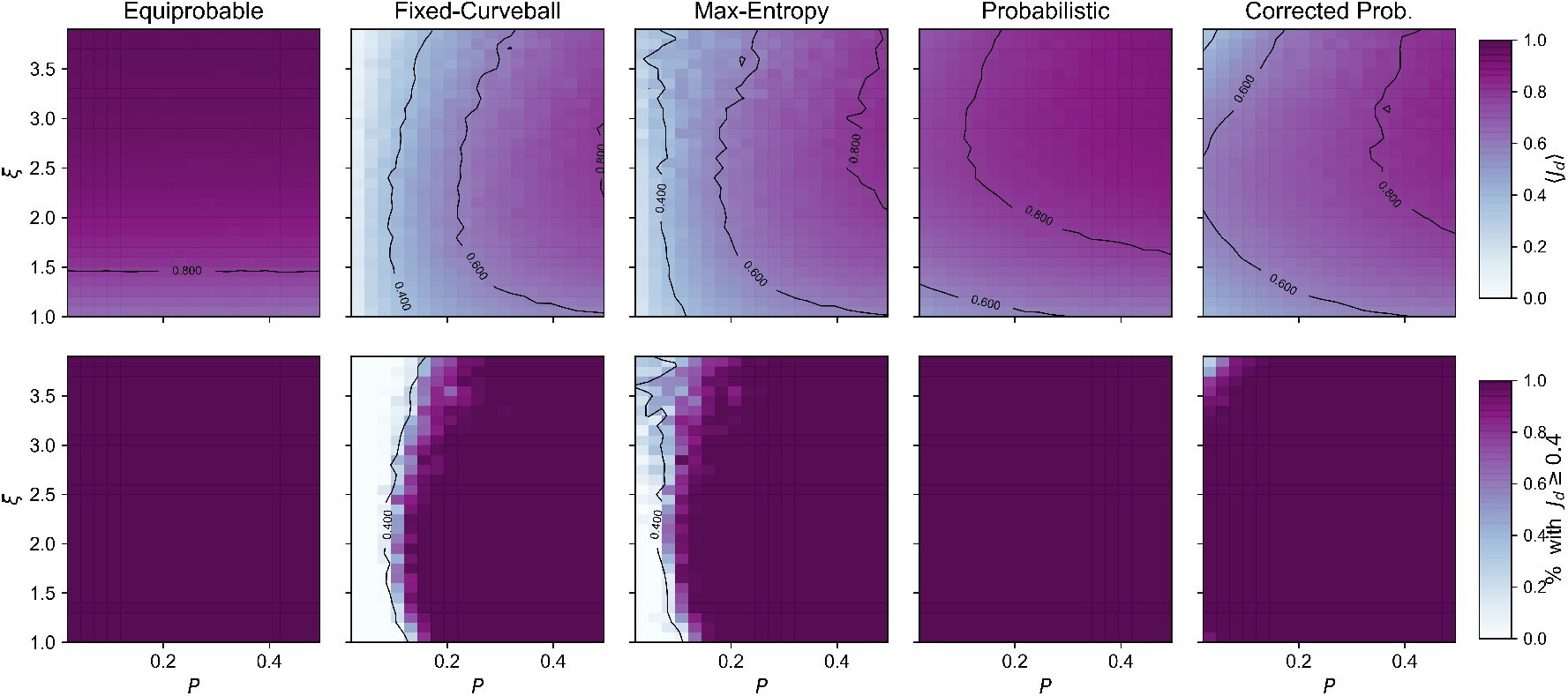
Synthetic nested Networks: 2 −dimensional plot in the *ξ* −*p* parameter space showing: the average Jaccard distance ⟨*J*_*d*_⟩ with of the null model ensembles with respect to the original nested synthetic matrix (top panel), and the % of matrices in the null model ensembles for which the Jaccard distance with respect to the original synthetic matrix was above a threshold of *J*_*d*_ ≥ 0.4 (bottom panel), for the five models studied in this work.

From another perspective, the bottom panel of Fig. 4 shows the percentage of matrices in the null model ensembles for which *J*_*d*_ ≥ 0.4 in the nested *ξ*− *p* parameter space. Clearly, there are relevant regions for which the FF and the Max-entropy models cannot generate true variants of the original matrix, even with such moderate threshold. Of course, when imposing a more rigid threshold this region becomes wider. In the other extreme, the EE and Probabilistic schemes prove to have ample room to generate largely different networks, no matter the parameter values under study. Somewhere in between, the Corrected Probabilistic Model is flexible enough to produce different networks, except in some extreme cases (very sparse networks with low *p*). These insights are all in good agreement with the work by Strona *et al*. [17], in which they proved that slightly relaxing the restrictiveness in the preservation of the rows and column totals –quasi-FF matrices– can reduce the tendency to under-estimate significant nestedness.

### B. Statistical significance of structural patterns

We now analyze how consistent the five null models described above are in assessing the statistical significance of nested, modular and in-block nested patterns, for each set of the generated synthetic matrices (192 empirical ecological networks are analyzed in the next section). In each case, we assessed the significance of the three patterns of interest by means of *z*-scores, as obtained from the average and standard deviations of an ensemble of randomized networks. The results of the analysis are summarized in Table I, which accounts the percentage of networks with *z* ≥ 2, *i*.*e*., an associated *α/*2 confidence limit around 0.025. For nested networks, we also explore the behaviour of the null models under three nestedness metrics: 𝒩 [24], *NODF* [51] and the spectral radius 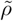 [13, 52], normalized with respect to the square root of the number of links in the matrices, 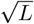, as in [53].

**TABLE I.** Percentage of networks with *z* ≥ 2 for three sets of synthetic networks with nested, modular and in-block nested structure. The over- and underestimation values mentioned in the text are marked in red and blue, respectively.

| Planted structure |  | EE | FF | PP |  |  |
| --- | --- | --- | --- | --- | --- | --- |
|  |  | Equiprobable | Fixed Curveball | Max-entropy | Probabilistic | Prob. Corrected |
| Nested | $\mathcal{N}$ | 100* | 1.28** | 5.9** | 99.88* | 41.33 |
| | $NODF$ | 100* | 0.06** | 4.73** | 99.93* | 46.21 |
| | $\tilde{\rho}$ | 100* | 28.13** | 3.48** | 99.02* | 58.39 |
| | $Q$ | 6.24 | 24.2 | 46.37* | 11.28 | 19.32 |
| | $\mathcal{I}$ | 87.53* | 4.04** | 7.33** | 85.84* | 44.65 |
| Modular | $\mathcal{N}$ | 0.76 | 1.63 | 0. | 0. | 0. |
| | $Q$ | 72.96 | 72.34 | 85.99 | 70.74 | 81.12 |
| | $\mathcal{I}$ | 50.93 | 52.94 | 40.21 | 44.57 | 38.63 |
| In-block nested | $\mathcal{N}$ | 74.81* | 32.76* | 0 | 28.28* | 0.05 |
| | $Q$ | 56.82 | 87.64 | 97.1 | 84.71 | 92.86 |
| | $\mathcal{I}$ | 55.89 | 69.75 | 67.99 | 69.05 | 63.22 |

\* Type I error
\*\* Type II error

For networks with planted nested structure (*B* = 1, *p* ∈ [0, 0.5]), the EE and Probabilistic models overestimate systematically the statistical significance for nestedness –no matter the metric. In the other extreme, both the FF and Max-entropy restrictive schemes deliver an underestimation of the amount of truly nested networks. Thus, confirming their proneness to Type I and II errors, respectively. Only the Corrected Probabilistic model renders a balanced assessment, consistent with the parameters used in the generation of this set of nested synthetic matrices. The assessment of the significance for in-block nestedness is generally consistent with the one observed for nestedness, as ℐ reduces to 𝒩 when *B* = 1 (see [24]).

More intriguingly, the FF and Max-entropy models wrongly identify a large fraction of nested-by-construction networks as being significantly modular (24.2% and 46.37%, respectively). To some extent, this can be explained by the densest nested matrices in the generated collection [24]. Still, it can’t hardly make sense to identify a larger proportion of networks as modular than as nested, in a set with purposefully planted nested structure.

Turning to modular-by-construction networks (*B >* 1, *p* = 1, *µ* ∈ [0, 0.5]), we observe that –unlike the previous case– all models render intuitively correct results: these networks are unanimously assessed as non-nested, and mostly assessed as modular. Such result is not surprising, since by construction, all the networks present a clearly modular planted structure, and there is evidence that these two patterns may not be structurally compatible [54]. On the other hand, around half of the collection is assessed as being in-block nested. The latter consideration makes sense, noting that an in-block nested configuration implies modularity. That is, in-block nested matrices are always a subset of the more general modular category.

Finally, networks with planted in-block nestedness (*B* = 4, *p* ∈ [0, 0.5], *µ* ∈ [0, 0.5]) again arise inconsistencies among the randomization procedures. Some of them (EE, FF and Probabilistic) incorrectly assess a large amount of networks as being nested –this is particularly salient in the case of EE.

Considering the results in Table I, it is clear that nested structures are the most confusing given the outcomes of the different randomisation schemes. For this reason, we explore these inconsistencies in further detail in Figure 5 regarding only the nested-by-construction synthetic subset. The figure shows the *z*-score values for nestedness *z*_𝒩_ (top) and modularity *z*_*Q*_ (bottom) over the *ξ* − *p* parameter space of the generated nested matrices, considering the three PP models. Thumbnails of the resulting generated matrices over the parameter space are shown on the left part of the figure as a visual aid.

**FIG. 5.**
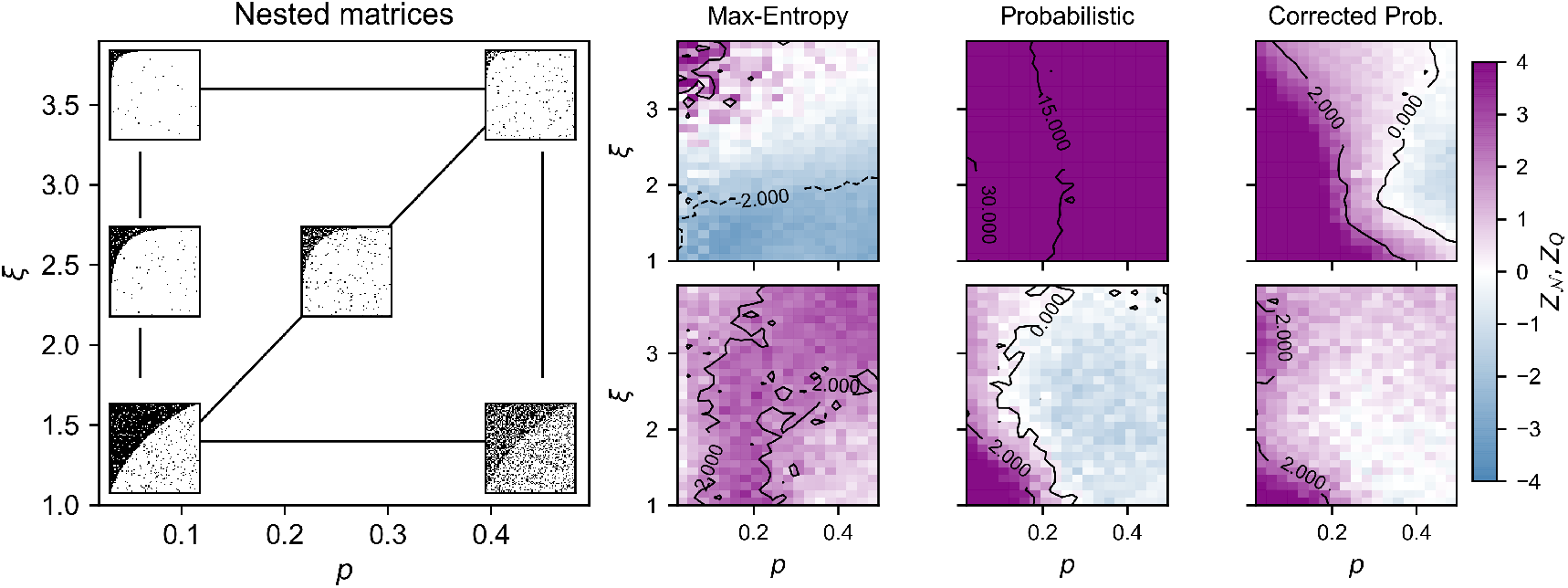
Synthetic Nested Networks: Heatmaps in the *ξ* − *p* parameter plane showing: *z*_*𝒩*_ (top) and *z*_*Q*_ (bottom) scores values under the Max-entropy, Probabilistic, and Corrected Probabilistic models for the set of ~ 9000 synthetic bipartite networks with planted nested structure. Examples of the resulting generated matrices over the parameter space are shown on the left part of the figure.

The results of Fig. 5 for the Probabilistic model confirm the tendency for overestimating the statistical significance of nested patterns, *i*.*e*., type I error. The model yields values of *z*_𝒩_ ≥ 2 over the whole parameter space, including the regions where the planted nested structure becomes hardly identifiable (high *p*). The Max-entropy model, conversely, is not able to assess the significance of nestedness, even in regions where the networks have a clearly defined nested pattern, except for those networks with high *ξ*. In fact, it deems a large amount of networks as, indeed, anti-nested, *i*.*e*., *z*_𝒩_ ≤ −2. The Corrected Probabilistic model is capable of assessing the significance of the nested pattern over a wide region of the parameter space, for intermediate to low values of noise *p* –as expected–, and different shape values *ξ*. Note, however, that the higher the value of *ξ* is, less noise *p* is necessary for the model to reach *z*_𝒩_ ≤ 2. This is not surprising, since the *ξ* parameter affects the network connectance, and sparser networks will require a lower level of noise to distort the planted nested structure. For the sake of completeness, Fig. S4 in the Supplemental Material shows the results for *NODF* and 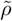, showing equivalent results, in spite of the different nature of the two metrics.

Focusing on the bottom row of Fig. 5, the assessment of significance for modularity *z*_*Q*_ with the Probabilistic and Corrected Probabilistic models shows small regions of *z*_*Q*_ ≥ 2 that correspond to high connectance networks (low *ξ*). In these cases, one can assume that networks are dense enough, such that the optimisation algorithm employed for modularity is able to find a partition with *B >* 1, resulting in values of *Q* larger than expected [24]. The Max-entropy model, however, confers significance for modular patterns over a wide region of the *ξ* − *p* parameter space –around 46% of the networks in Table I–, strengthening our assumptions regarding the presence of biases towards an overestimation of statistical significance of modular patterns.

Results for the assessment of modularity and in-block nestedness from Table I are complemented with Figure 6. The figure shows the *z*-score values for modularity *z*_*Q*_ (top) and in-block nestedness *z*_ℐ_ (bottom) over the *µ* − *B* parameter space of the generated matrices, with the three PP models. Similarly to Fig. 5, examples of the resulting matrices over the parameter space are shown on the left part of the figure. Again, Table I reports an unanimous assessment of modularity across the five models. Top panel in Fig. 6 for the PP models confirms these results: the models are able to assess the significance of modularity for a wide range of the parameter space. As expected, the assessment is sensitive to level inter-block noise *µ*. Results for the assessment of in-block nestedness also show consistent results across the five models (see Table I). For around half of the networks in this set, we obtain a significant level of in-block nestedness. Again, the critical ingredient here is the presence of inter-block noise *µ*. The result is in agreement with Palazzi *et al*. [54], in which it is proved that both *Q* and ℐ can coexist, as there are no mutually imposed constraints between them.

**FIG. 6.**
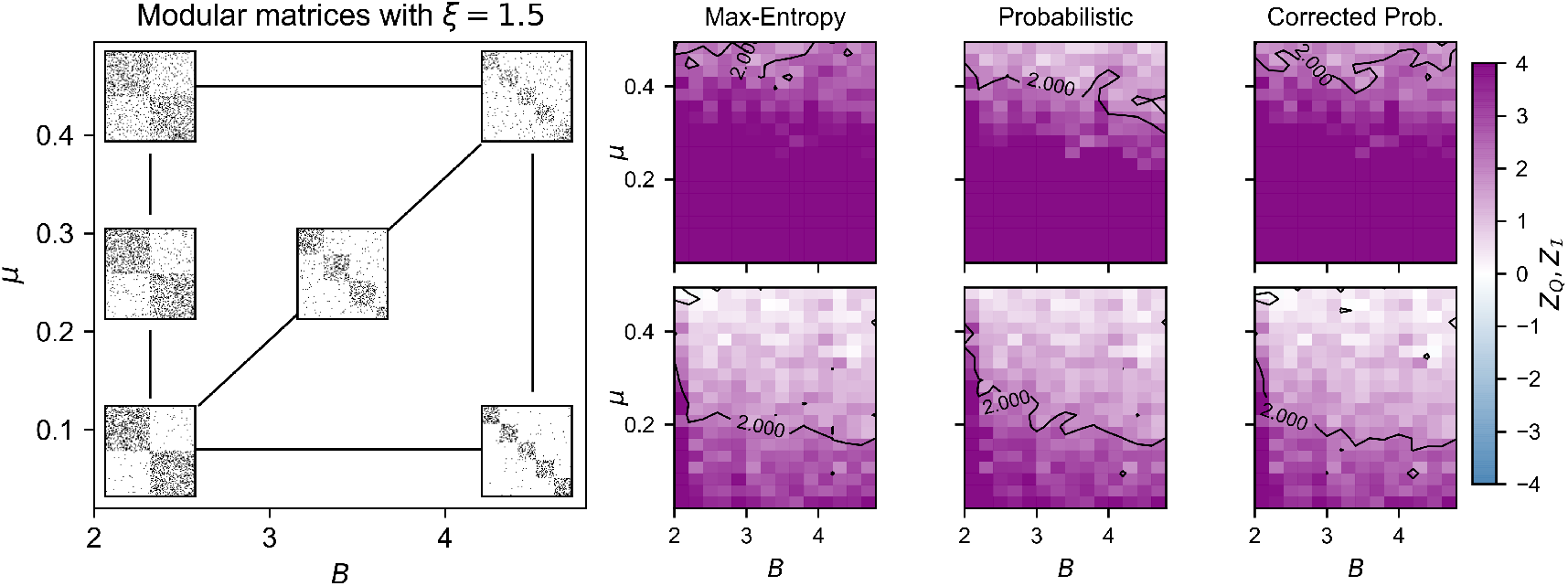
Synthetic Modular Networks: Heatmaps in the *µ* − *B* parameter plane showing: *z*_*Q*_ (top) and *z*_*ℐ*_ (bottom) scores values under the Max-entropy, Probabilistic, and Corrected Probabilistic models for the set of ~ 9000 synthetic bipartite networks with modular structure. Examples of the resulting generated matrices over the parameter space are shown on the left part of the figure.

Finally, we move on to examine the results for the last set of synthetic networks, with planted in-block nested structure. Figure 7 complements the results for this set of networks in Table I, by showing the *z*-score values for in-block nestedness *z*_ℐ_ (top) and nestedness *z*_𝒩_ (bottom). As in the previous cases, the left part of the figure shows examples of the generated matrices with the PP models over the *µ*−*p* parameter space. The three models illustrated here show similar (and expected) results for *z*_ℐ_: in-block nestedness becomes less significant when the structure is distorted within (higher *p*) or among (higher *µ*) blocks. The assessment of significance for 𝒩 in in-block nested networks (bottom row) is not as unanimous. In this case, it must be noted that global nestedness decreases with the number of blocks in a network, regardless of their internal nested structure [54]. Consequentially, the Max-entropy and the Corrected Probabilistic model correctly confer non-significant *z*_𝒩_ *<* 2 throughout the parameter plane, and even for low values of noise (either *µ* or *p*). Instead, the Probabilistic scheme identifies as nested those matrices for which *p <* 0.2, regardless of *µ*, a clear overestimation of nested patterns which occurs, even at higher proportions, in the EE and FF models (see Table I).

**FIG. 7.**
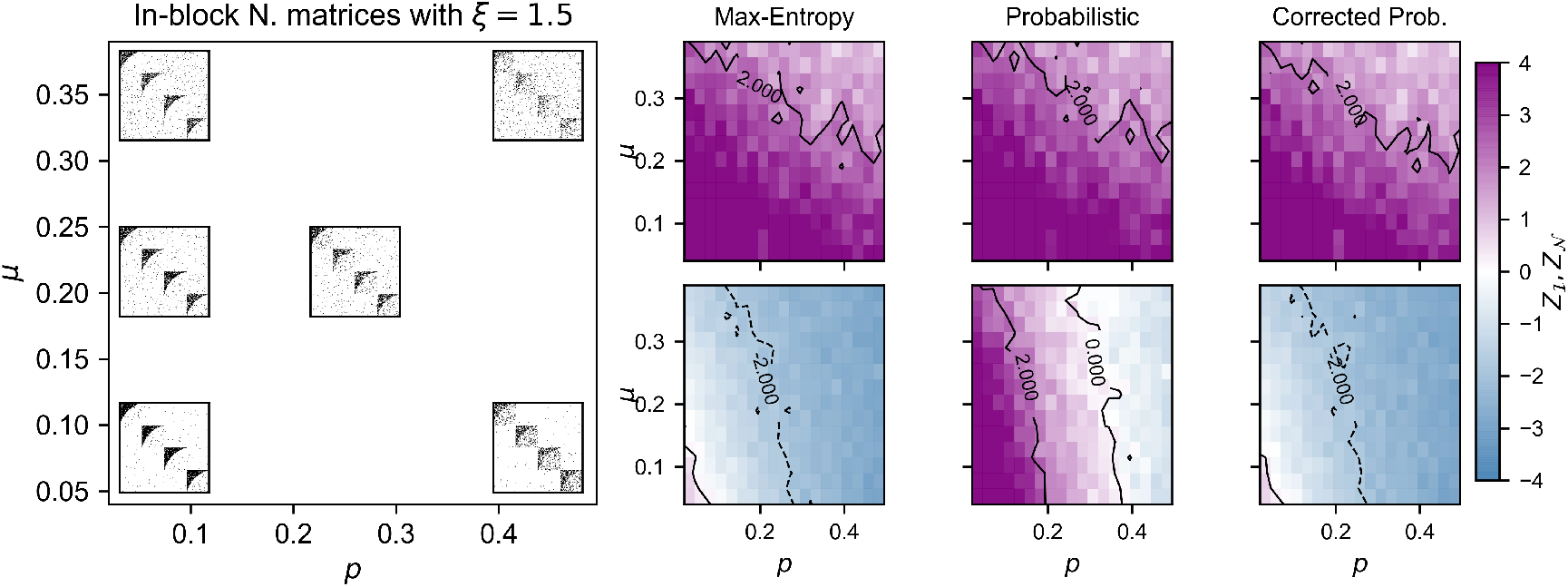
Synthetic in-block nested Networks: Heatmaps in the *µ* − *p* parameter plane showing: *z*_*ℐ*_ (top) and *z*_*𝒩*_ (bottom) scores values under the Max-entropy, Probabilistic, and Corrected Probabilistic models for the set of ~ 9000 synthetic bipartite networks with in-block nested structure. Examples of the resulting generated matrices over the parameter space are shown on the left part of the figure.

### C. Empirical Networks

~

Lastly, we analyze how consistent the five null models are in assessing the statistical significance of nested, modular, and in-block nested patterns over a set of 192 empirical networks [55]. All networks are treated as binary and have between 20 and 300 nodes (see Section S3 and Figure S5 in the Supplemental Material for details). For the sake of consistency and completeness, we explore the behaviour of the null models under the three nestedness metrics: 𝒩, NODF, and 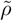. We report the percentage of networks with *z* ≥ 2 (*α/*2 ~ 0.025) for each one of our five structural metrics in Table II.

**TABLE II.**
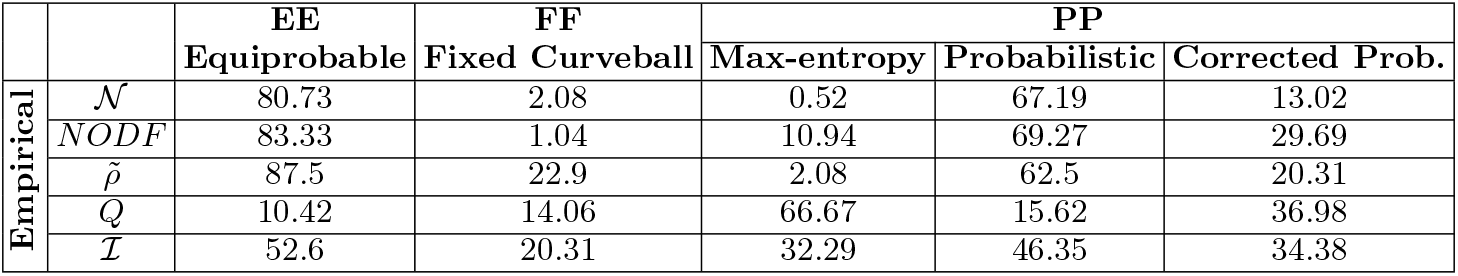
Percentage of networks with score *z* ≥ 2 for a collection of 192 empirical ecological networks.

Assessment of nestedness in the empirical data for the EE and the Probabilistic model is consistent across the three different metrics, yielding *z*-score values *z*_𝒩_ ≥ 2 for a large proportion of the analyzed networks (80.73% and 67.19%, respectively). The assessment of significance for 𝒩 in-block nestedness renders equivalent results, showing significant *z*_ℐ_ ≥ 2 for a high percentage of the networks, although in a lower proportion with respect to nestedness (52.6% and 46.35%, respectively). In contrast, only a small fraction of the networks is significantly modular (*z*_*Q*_ ≥ 2).

Results for the strictest schemes –FF and Max-entropy models–, show, in the case of nestedness, a good agreement with the results in the synthetic matrices (and the literature [49]), yielding non-significant *z*-scores for a large majority of the networks. Note that, similar to the synthetic case, the normalized spectral radius 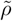, with the FF model is able to identify nestedness for a bigger proportion of the networks. Once again, the Corrected Probabilistic model seems to provide a more balanced assessment of nested patterns over the different metrics. The small variations in the preservation of the node’s degrees in the Corrected Probabilistic model allow nestedness to re-emerge as significant in a small proportion of the networks. Results for modularity in the FF and Max-entropy model render inconsistent results, suggesting once again the presence of biases in the Max-entropy model with regard to the assessment of modular patterns. Finally, we observe that a relevant fraction of the networks exhibit significant in-block nestedness structure even with the most restrictive algorithms, in close agreement with recent studies suggesting the presence of in-block nestedness as a frequent pattern in ecological communities [26, 28, 29].

## V. DISCUSSION

The overwhelming evidence that real ecological networks are highly structured, rather than randomly assembled, has sparked much research to explain the emergence and dynamical effects of those interaction patterns. In parallel, efforts have been devoted at determining the statistical significance of such patterns. As a result, the use and design of specialized null models to assess the statistical significance of selected architectures has become a widespread approach within the ecological community. However, to this day, there is still no agreement on which is the most suitable procedure to perform this kind of analysis. The discussion regarding the selection on the level of constraints to build the null matrices, the construction of synthetic benchmarks, or the appropriate null model-metric combination remains an open issue among scholars. Furthermore, there is already abundant evidence suggesting that networks often exhibit hybrid structures and/or at different scales of organisation, yet the concurrent assessment of the statistical significance of such complex patterns has not been extensively studied.

In this work, we have first addressed the need to build synthetic networks with total control over their dominant structure. Our generative benchmark allows for the construction of networks with clear defined architectural patterns, transitioning from structured (nested, modular, in-block nested) to random networks in a smooth way. Only then we can generate a rich sample to systematically study of the performance of five null models.

With over 25,000 networks of varying size, connectance and planted internal structure, we have observed the existence of ambiguities and biases in the concurrent assessment of network patterns, either over- or underestimating their significance. In particular, our use of an extensive ensemble of networks uncovers the weaknesses of both too restrictive (FF, Max-entropy) and too permissive (EE, Probabilistic) randomizing procedures. At the face of such results, we propose a variation of the Probabilistic model, that mitigates some of the limitations exhibited by the current models –in close agreement with recent research, regarding the convenience of quasi-FF approaches [17].

Our analysis of a rich synthetic ensemble shed some light on the reasons why the literature has reported contradicting results regarding the abundance of nested and/or modular networks in ecological systems. Our re-analysis of empirical networks under popular null model schemes reproduce those contradictions. Indeed, we observe again the presence of ambiguities in the simultaneous assessment of the patterns across models, with the Corrected Probabilistic model standing out as a balanced alternative to assess the significance of highly structured matrices.

We are convinced that our results can orient the future debate regarding the use of null models in bipartite networks within and beyond the ecological community. Providing a compact framework and off-the-shelf implementations (synthetic benchmark, structural descriptors, optimisation algorithms, and null models), our aim is double: on the theoretical side, to continue the research on the underlying assumptions and implicit consequences of the different null models. For example, to discern the apparent bias of the Max-entropy model towards underestimating the presence of nested configurations, while overestimating the presence of modular ones.

On the practical side, we seek to encourage researchers to shift the focus towards the simultaneous assessment of the statistical significance of multiple structural patterns at multiple scales –especially given the increasing evidence that empirical systems may display structure at several scales of organisation, and frequently in the form of compound structures. This is particularly crucial in ecology, in which transcending the traditional “nestedness vs. modularity” dichotomy marks a shift from the dominant approach in the mathematical modeling of ecosystems, and provides a consistent benchmark.

## DATA AND CODE AVAILABILITY STATEMENT

The software package BunGen for the generation of synthetic networks as described in Section II is freely available under the open source MIT licence. A stable version is available from https://github.com/COSIN3-UOC/BUNGen. The software package implementing the null models described in Section III (including the Corrected Probabilistic Model) is available from https://github.com/COSIN3-UOC/null_models. The data that support the results in Section IVC are openly available [55].

## ACKNOWLEDGMENTS

AL, AS-R and JB-H acknowledge financial support from the Spanish Ministerio de Ciencia e Innovación, through project No. PID2021-128966NB-I00. J.B-H. acknowledges financial support from the Ramón y Cajal program through the grant RYC2020-030609-I. AL also acknowledges financial support from the Spanish Ministerio de Ciencia e Innovación, through project No. PID2022-141802NB-I00 (BASIC).

