## Supplementary material for "Bipartite structural evaluation: Extended network generation model and corrected randomization techniques"

Harry de los Ríos[1], María J. Palazzi[1], Aniello Lampo[2], Albert Solé-Ribalta[1], Javier Borge-Holthoefer[1]

[1]Internet Interdisciplinary Institute (IN3), Universitat Oberta de Catalunya, Barcelona, Catalonia, Spain

[2]Grupo Interdisciplinar de Sistemas Complejos (GISC), Departamento de Matemáticas, Universidad Carlos III de Madrid, Spain

### S1. STRUCTURAL PATTERNS IN BIPARTITE NETWORKS: OVERVIEW

We present here the three structural arrangements considered in the manuscript: nestedness at the macroscale, modularity and in-block nestedness at the mesoscale. We provide their definition and discuss the related properties.

#### A. Nestedness

Nestedness, originally conceived in the field of biogeography [58–60], constitutes a frequent architectural pattern both inside [1, 6, 61] and outside Ecology [14, 62–65]. A perfect nested pattern describes a hierarchical organisation where the set of neighbours of a node is a subset of the neighbourhoods of larger degree nodes. That is, for the set of vertices $V$ in a graph, a perfectly nested structure exists if

$$\forall i, j \in V, \quad k_i < k_j \iff \Gamma_i \subset \Gamma_j, \tag{S1}$$

where $k_i$ and $\Gamma_i$ indicate the degree of a node $i$ and its neighbourhood, respectively. In the case of bipartite networks, this definition is valid provided one compares nodes belonging to the same class.

In terms of adjacency matrices, this definition translates into the characteristic triangular shape, as that shown in Figure S1 (left). In terms of visualisation, after re-ordering matrix rows and columns by degree, there exists a monotonic separatrix above which all the elements are greater than zero (or simply 1, in the case of presence-absence networks), and zero below. Here, nodes with the highest degree represent generalist species, while those with the lowest degree constitute the specialist ones. The form of this separatrix, defining the nested profile, enables many network configurations grouped under the umbrella of nestedness. This expresses the mutual proportion between generalists and specialists: tight separatrices describe communities with few generalists and a large number of specialists with a low number of interactions, while uniform profiles refer to a more balanced distributions of generalists and specialists. Noteworthy, perfectly nested patterns are extremely rare in real ecosystems. Even those networks exhibiting a highly nested character present links below the separatrix.

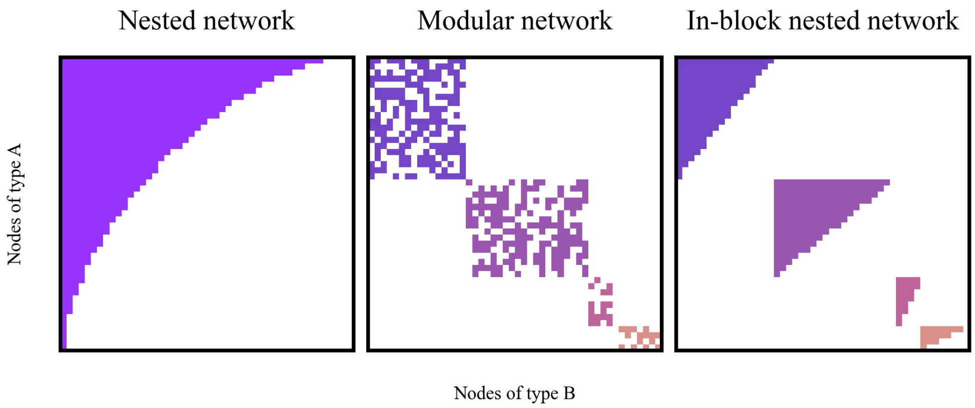


FIG. S1. Idealised examples of adjacency matrices associated to nested (left), modular (middle), and in-block nested (right) networks. For perfectly nested networks, the adjacency matrix manifests a triangular structure, while in the modular case it is divided in blocks, exhibiting a high internal connectivity (otherwise unstructured) and a low inter-block one (in this idealised example, none). The adjacency matrix of an in-block nested matrix is divided in blocks with an internal nested structure.

So far, this rich variety of nested networks has been characterised by introducing a large amount of metrics [66]. From an algebraic perspective, the spectral properties of perfectly nested graphs [52, 67] facilitated the proposal of Staniczenko *et al.* [13], which quantifies nestedness with respect to the maximum eigenvalue of adjacency matrices. Besides older approaches (e.g., Nestedness Temperature [68]), overlap-based measures are the most popular way to evaluate nestedness. Among those, the Node Overlap and Decreasing Fill (NODF) [51] stands out, along with its variants –weighted [69–71] or overlap-expectation-corrected [24] versions.

#### B. Modularity

Modular networks are those built up from weakly interlinked groups (also called communities, modules, compartments, or clusters) with high internal connectivity. Figure S1 (middle) depicts the adjacency matrix of a perfectly modular network, *i.e.* with no intergroup links. Modular structure is a rather ubiquitous mesoscale architecture [72] and appears in several fields, as diverse as biology [73] and cognitive science [74], including of course ecology [2, 23, 25, 75, 76].

The problem of identifying community structure constitutes itself a sub-area of network science. Over the last decades, scholars have developed a rich collection of algorithms and methodologies to infer these communities from relational data [72, 77]. One of the most popular methods in network science, and in particular in ecology, is the maximisation of modularity $Q \in [0, 1)$ [78, 79], which has been implemented in a myriad of optimization strategies [77, 80]. Of interest in this work, we focus on Barber's [79] formulation of modularity $Q$ for bipartite networks,

$$Q = \frac{1}{L} \sum_{i=1}^{N} \sum_{j=N+1}^{N+M} \left( \tilde{b}_{ij} - \tilde{p}_{ij} \right) \delta(\alpha_i, \alpha_j) \tag{S2}$$

where $L$ is the number of interactions (links) in the network, $\tilde{b}_{ij}$ denotes the existence of a link between nodes $i$ and $j$, $\tilde{p}_{ij} = k_i k_j / L$ is the probability that a link $(i, j)$ exists by chance, and $\delta(\alpha_i, \alpha_j)$ is the Kronecker delta function, which takes the value 1 if nodes $i$ and $j$ are in the same community, and 0 otherwise.

It is worth highlighting that current trends in pattern identification are increasingly skeptical about heuristic-based methods, and suggest the need to turn to better grounded methods like Bayesian inference –a demand that the synthetic generator's probabilistic formulation, as proposed in this work, can easily fulfil.

Worth noting, a modular architecture does not presuppose any particular in-block organisation: modularity implies the existence of groups in a network, but connectivity within those groups is not specified –it is implicitly assumed to be random. With exceptions for unipartite networks only [81, 82], a suitable benchmark to implement this architectural pattern by controlling the related ecology-relevant parameters is still missing.

#### C. In-block nestedness

Theoretical and empirical contributions in the last decade suggest that many complex systems may exhibit jointly nestedness and modularity. The existence of concurrent nested and modular organisations has been debated in different situations [2, 65], and yet empirical [83] and later analytical [54] evidence proved that macroscale nestedness and modular mesoscale cannot coexist easily.

This apparent incompatibility is due to the fact that the two patterns arise as a consequence of different mechanisms: certain pressures promote the block organisation, while others favour the emergence of nested patterns. Actually, the analysis of these mechanisms suggests that the co-occurrence of nestedness and modularity may take place at different scales. Herein, in-block nestedness stands out as a hybrid architecture that results from recasting nestedness at the mesoscale, *i.e.*, within modules [23, 24, 28].

In-block nestedness depicts a network configuration where weakly connected blocks exhibit an internal nested assembly (see Figure S1, right panel), and was first introduced by Lewinsohn *et al.* [22]. Such pattern has been found to unfold from an abundance-maximisation process [4] on top of a niche-structured population [26, 27], providing a bottom-up perspective on the emergence of nested-modular architectures. Notably, in-block nested networks present a balance between stability and diversity in mutualistic communities [29], and in-block nestedness is a predominant pattern in a significant amount of real plant-animal communities [29], and beyond [84].

Current approaches to detect nested compartments use a specialized fitness function [24]. For bipartite networks, it takes the form:

$$\mathcal{I} = \frac{2}{N_r + N_c} \left\{ \sum_{s,t}^{N_r} \left[ \frac{O_{s,t} - \langle O_{s,t} \rangle}{k_t(C_s - 1)} \Theta(k_s - k_t) \delta(\alpha_s, \alpha_t) \right] + \sum_{\sigma,\tau}^{N_c} \left[ \frac{O_{\sigma,\tau} - \langle O_{\sigma,\tau} \rangle}{k_\tau(C_\sigma - 1)} \Theta(k_\sigma - k_\tau) \delta(\alpha_\sigma, \alpha_\tau) \right] \right\} \tag{S3}$$

where $s$ and $t$ correspond to nodes in one class, and $\sigma$ and $\tau$ to nodes in the other. $O_{\cdot,\cdot}$ measures the degree of overlap between row and column pairs, and $\langle O_{s,t} \rangle = k_s k_t / N_c$ the expected amount of shared neighbors, *i.e.*, the expected overlap between two nodes. Finally, $\Theta(\cdot)$ is the Heaviside step function (such that the only contributing terms are those in which the outer index has larger degree than the inner).

In-block nestedness detection, like modularity, is a hard computational problem where the use of heuristic algorithms is most frequent. In this sense, the strategies to identify in-block nested arrangements (that is, in-block nestedness maximization) are similar to (and suffer from the same shortcomings as) those used in modularity.

### S2. PROBABILISTIC MODELS

#### A. Probability matrices and degree sequences

Following the template of Fig. 3, Section III of the main text, Figure S2 shows a certain realization of a modular (a) and an in-block nested (b) network. To their right, we see the resulting interaction probability matrices, produced by the three PP models, and scatter plots showing the fit between the obtained average degree sequences and the real degrees. For the Probabilistic model, we observe that over- and under-estimation of lower and higher degrees, respectively, persist, as in the case of nested matrices (Fig. 3 in the main text). On the other hand, we observe that both the Max-entropy and the Corrected Probabilistic models successfully preserve the average degree sequences of the modular and in-block nested synthetic matrices. Note worthy, both models present more homogeneous probability distributions across the matrix, thus granting them ample flexibility to generate degree-preserving randomized matrices that are substantially different from the original ones.

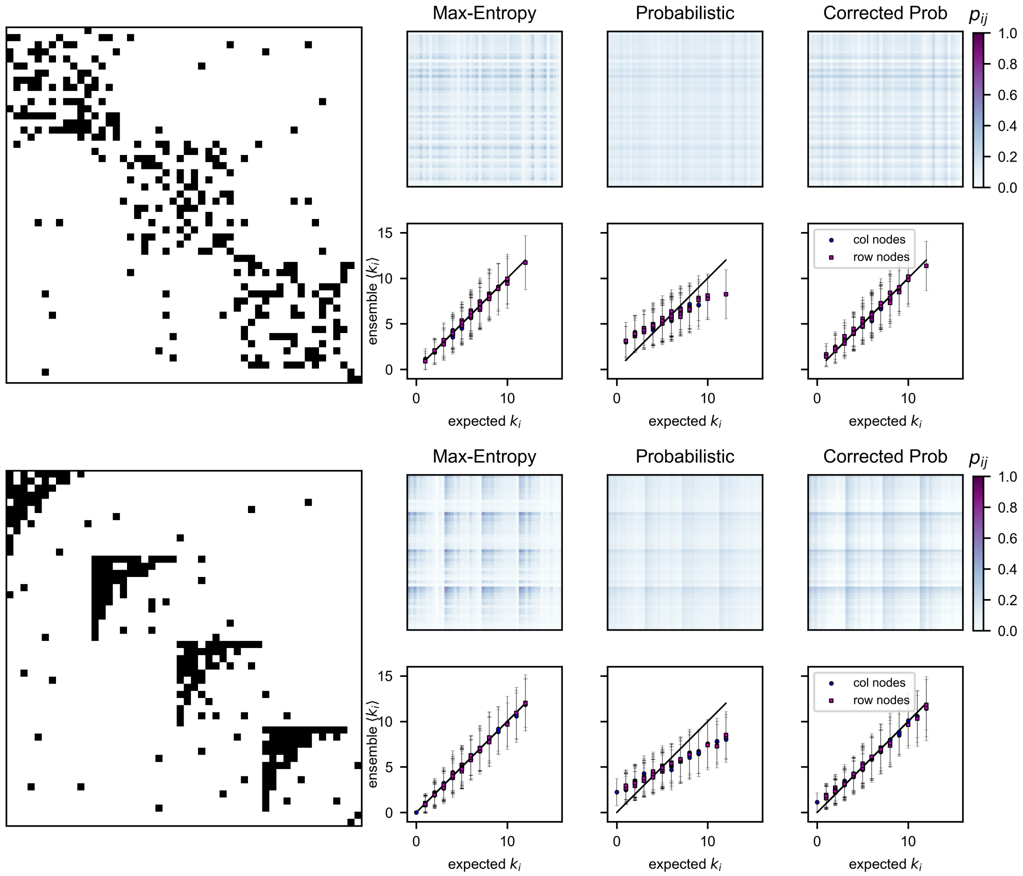


FIG. S2. Comparison between a synthetic network (left) with planted modular (a) and in-block nested (b) structures and their corresponding matrices of interaction probabilities $p_{ij}$ produced by the three PP models: maximum-entropy, Probabilistic and Probabilistic Corrected, respectively, and the scatter plots showing the fit between the obtained average degree sequences with each model and the real degrees of the corresponding matrix. As in Fig. 3 of the main text, error bars in the scatter plots indicate one standard deviation above and below the average.

### B. Original vs randomized networks: Jaccard Distance

In this section, in a similar manner of Fig. 4, Section IV A of the main text, we take a closer look at the ensemble of null matrices generated by each one of the three models over the parameter space of our benchmark graph model, for the modular and in-block nested synthetic networks. We explore to what extent the generated null matrices resemble the original synthetic ones for the whole range of parameters of the graph model, by computing the Jaccard distance $J_d$.

Top panel in Figure S3 shows the average Jaccard distance $\langle J_d \rangle$ over the modular $\mu - B$ parameter space for each PP model. The bottom panel corresponds to the in-block nested $\mu - p$ parameter space. Free from the tight restrictions of nested patterns, for all models we observe high values of the $\langle J_d \rangle$ over the $\mu - B$ and $\mu - p$ parameter spaces, respectively, showing that for the cases of networks with modular or in-block nested structure, the null matrices generated by these models are substantially different from the original matrix.

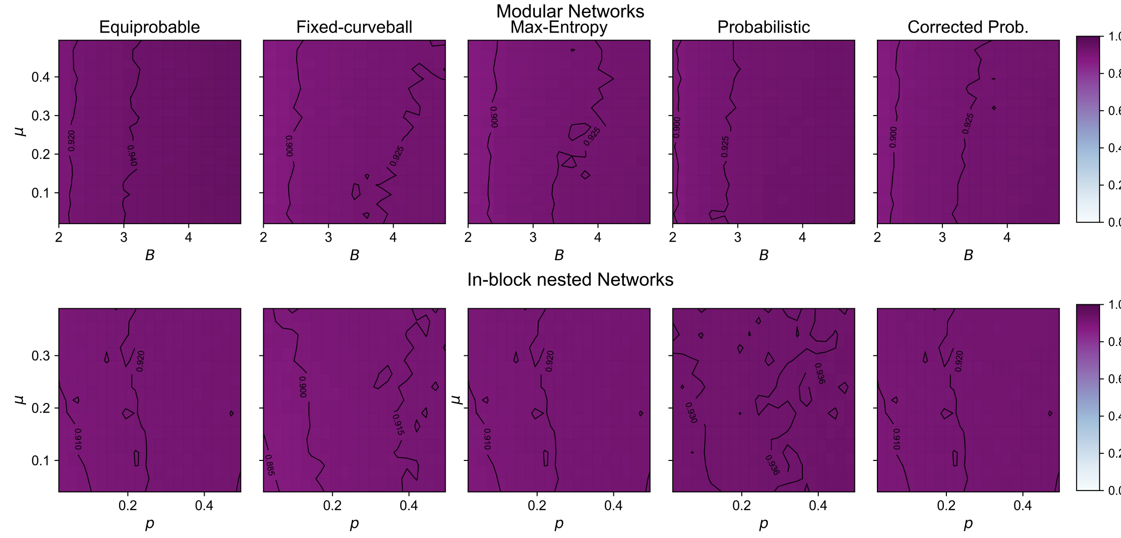


FIG. S3. 2−dimensional plot in the $\mu - p$ parameter space showing the average Jaccard distance $\langle J_d \rangle$ with of the null model ensmebles with respect to the original synthetic matrix (modular and in-block nested in the top and bottom rows, respectively) for the three types of PP models studied in this work.

#### C. Statistical significance of nestedness under different metrics: NODF and Spectral Radius)

In Section IV B of the main text, we have shown that the assessment of the statistical significance of nested patterns can lead to inconsistent and/or ambiguous results across different null models. To this aim, we employed the nestedness metric introduced by Solé-Ribalta *et al.* [24], an overlap metric that includes a term that discounts the expected overlap coming from random interactions. For the sake of completeness, in this section, we explore this behavior across null models under other nestedness metrics. Specifically, we employ the well-known NODF [51], and the spectral radius [52], normalized with respect to the number of links in the matrices, as in [53].

Figure S4 shows $z_{NODF}$ (top) and $z_{\tilde{\rho}}$ (bottom) $z$-scores values under the Max-entropy, probabilistic, and the corrected probabilistic null models in the $\xi - p$ parameter space, showing overall equivalent results from those presented in Fig. 6 (top panel) of the main text. Differences in the regions of transition from significant to no significant nestedness for $z_{\mathcal{N}}$ and $z_{NODF}$ in the $\xi - p$ space, despite of both being overlap metrics, are due to the effect of the null model term on the $\mathcal{N}$ metric.

### S3. DISTRIBUTION OF THE NUMBER OF SPECIES AND CONNECTANCE OF THE EMPIRICAL NETWORKS

Figure S5 shows the distribution of the parameters of connectance $C$ and the number of species $S$ for the empirical mutualistic networks which are analyzed in Section IV C of the main text [55]. From both the box plot and the histogram distributions of $S$ and $C$, we observe that almost the majority of the networks (at a 95%CI) show values of $S < 110$ and $C < 0.3$. Network sizes and connectance above these thresholds could be considered outliers. Hence, we restricted the generation of synthetic ensembles to networks within those values.

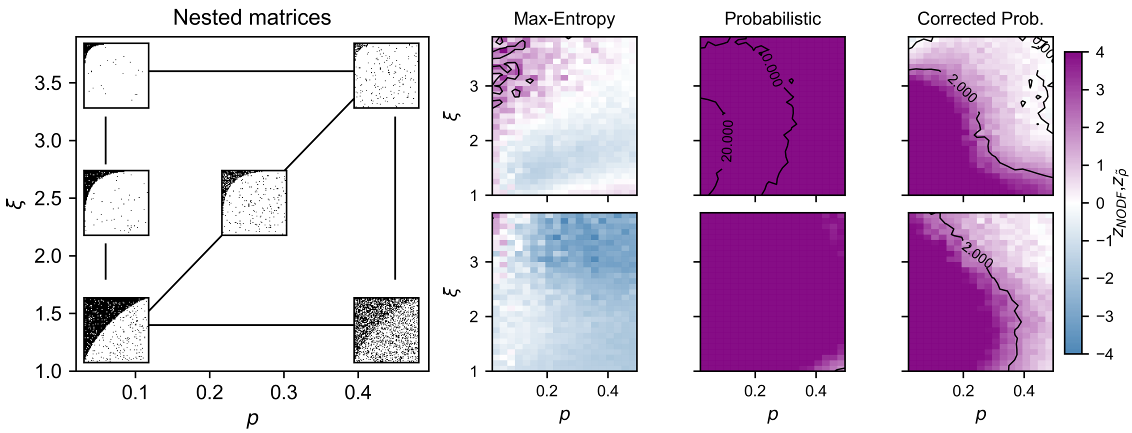


FIG. S4. **Synthetic Nested Networks:** 2−dimensional plots in the $\xi - p$ parameter space showing: $z_{NODF}$ (top) and $z_{\tilde{\rho}}$ (bottom) $z$-scores values under the entropy-based, the probabilistic, and the corrected probabilistic null models for the set of ∼9k synthetic bipartite networks with nested structure. Examples of the resulting generated matrices over the parameter space are shown on left part of the figure.

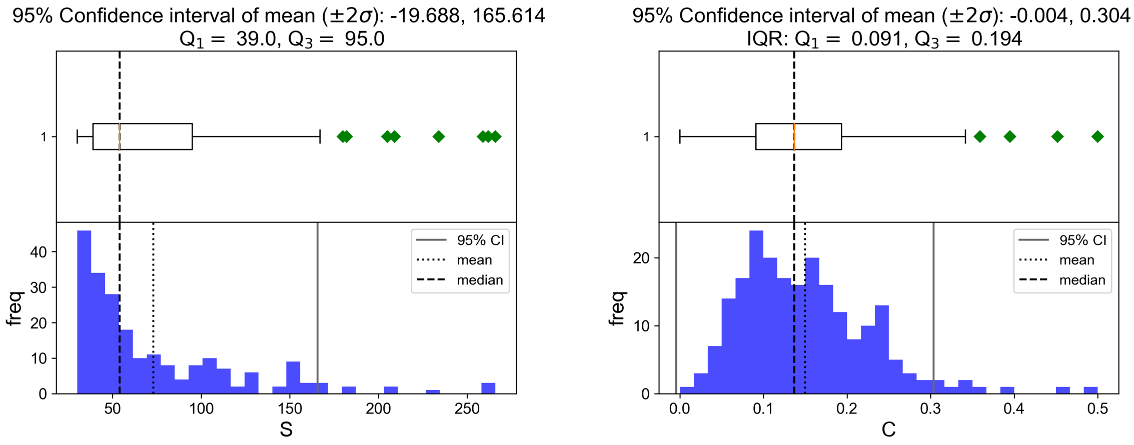


FIG. S5. Box plots (top) and histograms (bottom) showing the distributions of parameters $C$ (left panels) and and $S$ (right panel) of the real mutualistic networks. Dashed black lines in all panels indicates were the median value for each parameter is located, dotted black line on the histograms correspond to the mean, and solid purple lines show the values above or below the 95% confidence interval of the mean.

[32] T. P. Peixoto, *Descriptive vs. inferential community detection in networks: Pitfalls, myths and half-truths* (Cambridge University Press, 2023).

.

[56] BUNGen: Synthetic generator for structured ecological networks, `https://github.com/COSIN3-UOC/BUNGen` (2024)

.

[57] Null models implementation, `https://github.com/COSIN3-UOC/null_models` (2024)

.

[58] E. Hultén, *Outline of the history of arctic and boreal biota during the Quaternary period* (Bokforlags Aktiebolaget Thule, 1937).

[59] P. J. Darlington, Zoogeography, The Geographical Distributions of Animals (1957).

[60] B. D. Patterson and W. Atmar, Nested subsets and the structure of insular mammalian faunas and archipelagos, Biological Journal of the Linnean Society **28**, 65 (1986).

[61] S. Cobo-López, V. K. Gupta, J. Sung, R. Guimerá, and M. Sales-Pardo, Stochastic block models reveal a robust nested pattern in healthy human gut microbiomes, PNAS Nexus

(2022).
